# Coordination between nucleotide excision repair and specialized polymerase DnaE2 action enables DNA damage survival in non-replicating bacteria

**DOI:** 10.1101/2021.02.15.431208

**Authors:** Asha Mary Joseph, Saheli Daw, Ismath Sadhir, Anjana Badrinarayanan

## Abstract

Translesion synthesis (TLS) is a highly conserved mutagenic DNA lesion tolerance pathway, which employs specialized, low-fidelity DNA polymerases to synthesize across lesions. Current models suggest that activity of these polymerases is predominantly associated with ongoing replication, functioning either at or behind the replication fork. Here we provide evidence for DNA damage-dependent function of a specialized polymerase, DnaE2, in replication-independent conditions. We develop an assay to follow lesion repair in non-replicating *Caulobacter* and observe that components of the replication machinery localize on DNA in response to damage. These localizations persist in the absence of DnaE2 or if catalytic activity of the polymerase is mutated. Single-stranded DNA gaps for SSB binding and low-fidelity polymerase-mediated synthesis are generated by nucleotide excision repair, as replisome components fail to localize in its absence. This mechanism of gap-filling facilitates cell cycle restoration when cells are released into replication-permissive conditions. Thus, such cross-talk (between activity of NER and specialized polymerases in subsequent gap-filling) helps preserve genome integrity and enhances survival in a replication-independent manner.

## Introduction

DNA damage is a threat to genome integrity and can lead to perturbations to processes of replication and transcription. In all domains of life, bulky lesions such as those caused by UV light (cyclobutane pyrimidine dimers, CPD and to a lesser extent 6,4 photoproducts, 6-4PP) are predominantly repaired by Nucleotide Excision Repair (NER) (Boyce & Howard-Flanders, 1964; Chatterjee & Walker, 2017; Kisker et al., 2013). This pathway can function in global genomic repair (GGR) via surveilling the DNA double-helix for distortions or more specifically via transcription-coupled repair (TCR) (Kisker et al., 2013). The main steps of NER involve lesion detection followed by incision of few bases upstream and downstream of the lesion, resulting in removal of a short stretch of single-stranded DNA (ssDNA). This ssDNA gap is then filled by synthesis from a DNA polymerase (Kisker et al., 2013; Sancar & Rupp, 1983). While the NER-mediated damage removal pathway is largely error-free, lesions encountered by the replication machinery (for example CPDs, 6-4PPs and crosslinks such as those generated by antibiotics including Mitomycin C (MMC)) can also be dealt with via error-prone translesion synthesis (TLS) (Chatterjee & Walker, 2017; Fuchs & Fujii, 2013; Fujii & Fuchs, 2004).

TLS employs low-fidelity polymerases to synthesize across DNA lesions, with increased likelihood of mutagenesis during this process (Fuchs & Fujii, 2013; Galhardo, 2005; Kato & Shinoura, 1977; Nohmi et al., 1988; Warner et al., 2010). In most bacteria, expression of these polymerases is regulated by the SOS response, which is activated by the RecA-nucleoprotein filament under DNA damage (Baharoglu & Mazel, 2014). Currently most of our understanding about TLS comes from studies on specialized Y-family polymerases of *E. coli*, DinB (PolIV) and UmuDC (PolV), both of which function in DNA lesion tolerance and contribute to mutagenesis in several bacterial systems (Kato & Shinoura, 1977; Nohmi et al., 1988; Steinborn, 1978; Sung et al., 2003; J. Wagner et al., 1999). In addition, PolV has also been implicated in RecA-dependent post-replicative gap-filling activity (Isogawa et al., 2018). In contrast to *E. coli, Caulobacter crescentus* as well as other bacteria including *Mycobacterium sp.* and *Pseudomonas sp.* encode an alternate, SOS-inducible error-prone polymerase, DnaE2 (Galhardo, 2005; Jatsenko et al., 2017; Warner et al., 2010). DnaE2 is highly conserved and thought to be mutually exclusive with UmuDC in occurrence. In the limited organisms where DnaE2 has been studied so far, it is the primary TLS polymerase and the only contributor to damage-induced mutagenesis (Alves et al., 2017; Galhardo, 2005; Warner et al., 2010). In contrast to PolV, DnaE2 is thought to preferentially act on MMC-induced damage, where it contributes to all induced-mutagenesis observed. In case of UV, there are still uncharacterized mechanisms that can contribute to damage tolerance and mutagenesis that are independent of DnaE2 (Galhardo, 2005). DnaE2, co-occurs with ImuB, a protein that carries a β-clamp binding motif, and is thought to act as a bridge between DnaE2 and the replisome (Warner et al., 2010). Unlike *E. coli,* where activities of PolIV and PolV are well-studied, *in vivo* investigations of DnaE2 function in damage tolerance and its contribution to cellular survival are limited. This becomes particularly important, given the emerging evidences across domains of life ascribing diverse functions to these low-fidelity polymerases beyond their canonical function of replication-associated lesion bypass (Joseph & Badrinarayanan, 2020). Indeed, such polymerases are also referred to as ‘specialized polymerases’ (Fujii & Fuchs, 2020) so as to consider these broader functions.

Since these error-prone polymerases can synthesize DNA and their activity is mediated by interaction with the β-clamp of the replisome (Bunting et al., 2003; Chang et al., 2019; Fujii & Fuchs, 2004; Thrall et al., 2017; Jérôme Wagner et al., 2009; Warner et al., 2010), action of these polymerases has mostly been studied in the context of replicating cells, as a mechanism that facilitates continued DNA synthesis by acting at or behind the replication fork (Chang et al., 2019, 2020; Indiani et al., 2005; Jeiranian et al., 2013; Marians, 2018). In addition to replication-associated lesion tolerance, some studies have proposed the possibility of error-prone synthesis in a manner that is replication-independent (Janel-Bintz et al., 2017; Kozmin & Jinks-Robertson, 2013). This is supported by observations that cells can undergo stationary phase mutagenesis that is dependent on action of error-prone polymerases (Bull et al., 2001; Corzett et al., 2013; Janel-Bintz et al., 2017; Sung et al., 2003; Yeiser et al., 2002). Microscopy-based approaches have also provided evidence in line with the idea that tolerance or gap-filling could occur outside the context of the replication fork in *E. coli,* as replisome components, such as the β-clamp, as well as specialized polymerases (PolIV and PolV) were found to localize away from the fork in response to DNA damage (Henrikus et al., 2018; Robinson et al., 2015; Soubry et al., 2019; Thrall et al., 2017). Furthermore, while originally considered as distinct mechanisms of repair (damage tolerance vs damage removal), recent studies also suggest cross-talk between specialized polymerases and NER in *E. coli,* yeast and human cells (Giannattasio et al., 2010; Janel-Bintz et al., 2017; Kozmin & Jinks-Robertson, 2013; Sertic et al., 2018). Indeed, long-standing observations suggest that NER can be mutagenic under certain conditions in *E. coli*, in a manner that is dependent on RecA (Bridges & Mottershead, 1971; Cohen-Fix & Livneh, 1994; Nishioka & Doudney, 1969). However, the mechanistic basis of this process in replication-independent conditions and conservation of the same across bacteria that encode diverse specialized polymerases remains to be elucidated. For example, unlike *E. coli*, several bacterial systems undergo non-overlapping cycles of DNA replication and have distinct cell cycle phases with no ongoing DNA synthesis. The relevance of lesion correction or gap-filling for genome integrity maintenance in the absence of an active replication fork (such as in non-replicating swarming cells) remains incompletely explored and more so in bacterial contexts.

To probe the *in vivo* mechanism and understand the impact of error-prone polymerase function in non-replicating bacteria, we investigated lesion repair in *Caulobacter crescentus* swarmer cells. *Caulobacter* is well-suited to study activity of these specialized polymerases due to its distinct cell cycle. Every cell division gives rise to two different cell types: a stalked and a swarmer cell. While the stalked cell initiates replication soon after division, a swarmer must differentiate into a stalked before replication re-initiation (Schrader & Shapiro, 2015) and hence swarmers represent a pool of naturally occurring non-replicating cells in the environment. Under laboratory conditions, these swarmer cells can be isolated via density-gradient centrifugation and replication initiation can be inhibited, resulting in a population of non-replicating cells with a single chromosome (Badrinarayanan et al., 2015; Schrader & Shapiro, 2015). Using this non-replicating system, we followed DNA damage repair with lesion-inducing agents via live-cell fluorescence microscopy. We show that low-fidelity polymerase DnaE2 is active and functional in gap-filling damaged DNA in non-replicating cells. This is facilitated by *de novo* loading of replisome components (SSB, HolB (part of the clamp loader complex), β-clamp and replicative polymerase) at ssDNA gaps generated by NER. We find that this form of gap-filling in non-replicating cells promotes cell cycle restoration and cell division, upon release into replication-permissive conditions. Our study provides *in vivo* evidence for a novel function of DnaE2 that is spatially and temporally separate from the active replication fork. Given that DNA damage can occur in any cell type whether actively replicating or not, coordinated activity of NER and low-fidelity polymerases can serve as a potential mechanism through which non-replicating cells such as bacteria in stationary phase or cells in other differentiated phases increase their chances of survival under damage.

## Results

### Monitoring mechanisms of DNA lesion repair in non-replicating bacteria

To test whether non-replicating cells can indeed engage in lesion repair, and understand the *in vivo* mechanism of such activity, we used *Caulobacter crescentus* swarmer cells as our model system. We regulated the state of replication so as to ensure that swarmer cells, with a single chromosome, do not initiate replication (and hence prevent possibility of recombination-based repair) by utilizing a previously described system to control the expression of the replication initiation protein, DnaA, from an IPTG inducible promoter (Badrinarayanan et al., 2015). In our experimental setup, we first depleted cells of DnaA for one generation of growth, followed by synchronization to isolate non-replicating swarmer cells (Figure 1A, top panel). Flow cytometry profiles of cells confirmed the presence of a single chromosome during the course of the entire experiment (Figure 1A, bottom panel).

**Figure 1:**
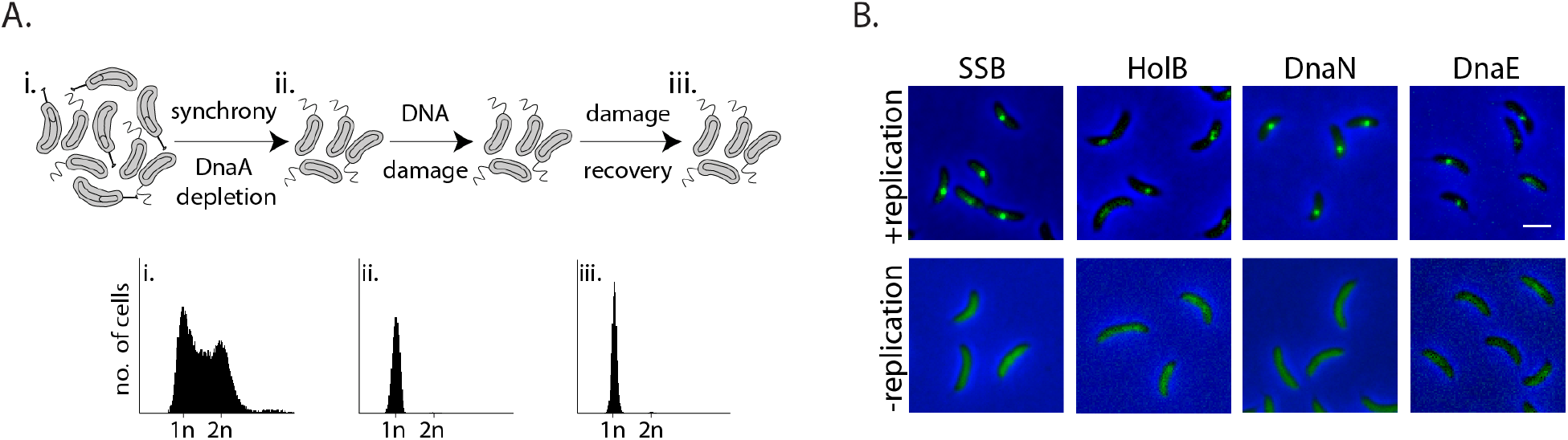
Monitoring mechanisms of DNA lesion repair in non-replicating bacteria. (A) above: Schematic of experimental setup used to isolate non-replicating *Caulobacter* swarmer cells to monitor DNA lesion repair and tolerance independent of ongoing replication. Cells are treated with DNA damage (30 min MMC or UV), after which damage is removed and cells are allowed to grow in fresh media (damage recovery), without ongoing replication. below: Flow cytometry profiles show DNA content in an asynchronous population (i), synchronized non-replicating swarmer cells before (ii) and after DNA damage recovery (iii). (B) Representative images of *Caulobacter* cells with fluorescently-tagged replisome components (SSB-YFP, HolB-YFP, DnaN-YFP or DnaE-mNeonGreen) in replicating or non-replicating conditions, without DNA damage (scale bar is 2 μm here and in all other images).

Given the requirement of the β-clamp for activity of specialized polymerases and evidence for damage-dependent changes in localization of replisome components such as SSB in actively replicating *E. coli* (Chang et al., 2019; Henrikus et al., 2018; Soubry et al., 2019; Thrall et al., 2017), we generated fluorescent fusions to the *Caulobacter* β-clamp (DnaN), component of the clamp loader complex (HolB), the replicative polymerase PolIII (DnaE), and single-strand binding protein (SSB), (using previously described approaches in *Caulobacter* (Aakre et al., 2013; Collier & Shapiro, 2009); and materials and methods) in order to visualize them in non-replicating swarmers. These fusions did not perturb the function of the proteins as cells displayed wild type growth dynamics in steady-state conditions (Figure S1A and S1B ‘control’). They also did not have increased sensitivity to DNA damage treatment via MMC or UV (Figure S1A & S1B). The fusion proteins localized on DNA in actively replicating cells (Figure 1B, +replication) and as anticipated, their localizations gradually shifted from one pole to the other within one cycle of DNA replication (Figure S1C). These observations are in line with previous reports of replisome dynamics in several bacterial systems including *Caulobacter crescentus, Bacillus subtilis* and *E. coli* (Aakre et al., 2013; Collier & Shapiro, 2009; Jensen et al., 2001; Lemon & Grossman, 1998; Mangiameli et al., 2017; Reyes-Lamothe et al., 2008). In contrast to actively replicating cells, replication-inhibited swarmer cells were devoid of replisome foci (Figure 1B), consistent with the idea that the localization of replisome components is indicative of active DNA replication.

### Replisome components are recruited to damaged DNA in non-replicating *Caulobacter* swarmer cells

Using the above described system, we treated non-replicating *Caulobacter* swarmer cells with Mitomycin C (MMC) to induce DNA lesions and followed DNA damage recovery via live-cell imaging to track dynamics of the β-clamp and other replisome components (Figure 1A). MMC is a naturally produced antibiotic that acts predominantly on the guanine residue of DNA, making three major forms of damage: mono-adducts, intra-strand crosslinks and inter-strand crosslinks (Bargonetti et al., 2010). In case of *Caulobacter,* it is thought that DnaE2 preferentially acts on MMC-induced damage as all mutagenesis associated with MMC treatment is mediated via action of this specialized polymerase; in absence of the polymerase, cells show high sensitivity to MMC treatment. To determine the range of MMC concentration for this study, we first assessed the viable cell count for a steady state population of wild type and *dnaE2* deleted cells across increasing concentrations of MMC treatment (0.125 μg/ml - 2 μg/ml) and focused on a treatment range where DnaE2 essentiality was observed (Figure S2A) and TLS-dependent mutagenesis has previously been reported (Galhardo, 2005).

We then went ahead and treated non-replicating swarmer cells with specified doses of MMC. We found that DNA damage treatment resulted in the formation of β-clamp foci in non-replicating cells (Figure 2A-B). This was found to be the case for other replisome components as well (Figure 2A-B). The percentage of cells with damage-induced β-clamp foci increased with increasing doses of MMC; at 0.125 μg/ml MMC 9% cells had β-clamp foci, while at higher doses of 0.75 μg/ml, foci were observed in 59% cells (Fig. S4C). To further characterize the dynamics of these localizations during the course of damage recovery, we released MMC-treated non-replicating swarmers into fresh media without damage and followed the localization of replisome components over time, but maintained the block on replication initiation, thus ensuring that cells carried only a single non-replicating chromosome during the course of the entire experiment (Figure 1A). Consistent with the possibility of dissociation during recovery, we found that percentage cells with DnaN localizations gradually decreased with time (Figure 2C) and across all doses of damage tested (Figure S4C). For example, after 30 min of 0.5 μg/ml MMC treatment, 52% cells on average had DnaN localization and at 90 min after damage removal, the number reduced to 30%. This pattern of localization after damage treatment, followed by reduction in percentage cells with foci during recovery was also observed in the case of SSB, HolB and DnaE (Figure 2D). Interestingly, we noticed that cells had more SSB localizations on average than DnaN. 14% cells had ≥2 DnaN foci after MMC treatment, while 37% cells harboured ≥2 SSB localizations, and this number dropped with increasing time in recovery (Figure 2D). Assessment of the extent of colocalization between DnaN and SSB further showed that 90% of DnaN foci colocalized with SSB (with distance of a DnaN focus from the nearest SSB localization being within 300 nm), while only 51% of SSB foci colocalized with DnaN (Figure S2B and S2C), suggesting that not all SSB may be associated with the β-clamp or that SSB could precede β-clamp localization.

**Figure 2:**
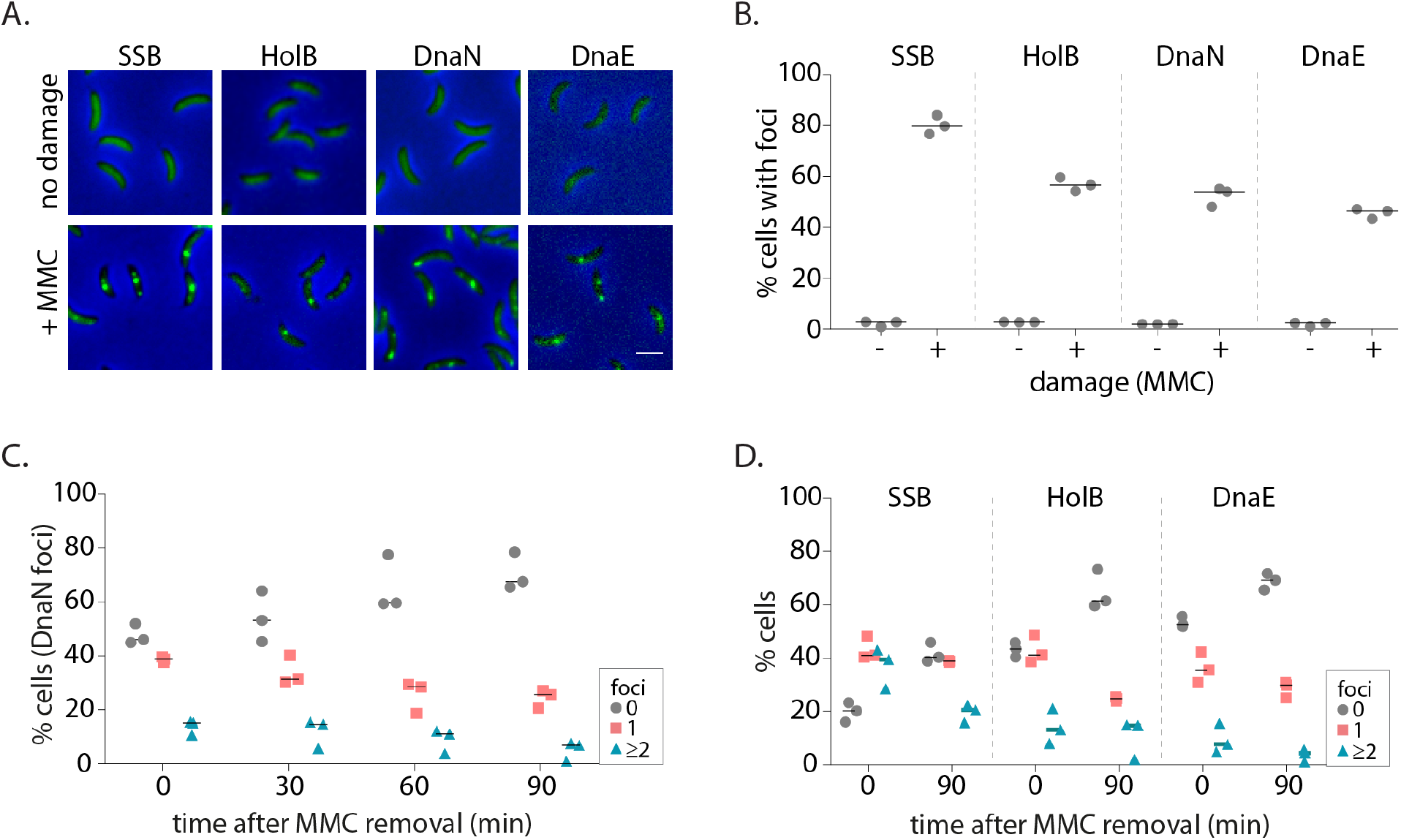
Replisome components are recruited to damaged DNA in non-replicating *Caulobacter* swarmer cells. (A) Representative images of non-replicating swarmer cells with fluorescently-tagged replisome components (SSB-YFP, HolB-YFP, DnaN-YFP or DnaE-mNeonGreen) with (+MMC) or without (no damage) 30 min of treatment with MMC. (B) Percentage cells with SSB, HolB, DnaN or DnaE localization (foci) in non-replicating swarmers with (+) or without (-) MMC treatment (n ≥ 324 cells, three independent repeats). Dashed line represents median here and in all other graphs. (C) Percentage swarmer cells with 0, 1, or ≥2 DnaN foci at 0, 30, 60 and 90 min after damage removal (recovery) (n ≥ 476 cells, three independent repeats). (D) Percentage swarmer cells with 0, 1, or ≥2 foci of SSB, HolB or DnaE at 0 and 90 min after damage removal (recovery) (n ≥ 324 cells, three independent repeats).

In order to further support observations made with MMC treatment, we asked whether these dynamics of replication machinery components were observed for another lesion-inducing agent as well. For this, we treated cells with sub-inhibitory doses of UV radiation that have been shown to have similar growth effects on wild type cells as MMC-treated *Caulobacter* (Galhardo, 2005 and Figure S2D). Exposure of cells to two doses of UV damage (75 J/m^2^ and 150 J/m^2^) also resulted localization and subsequent reduction in percentage cells with replisome foci during recovery (Figure S2E, S2F, S2G). Taken together, these data support the idea that SSB, along with components of the PolIIIHE, including the clamp-loader, β-clamp and the replicative polymerase, associate with DNA during damage even in the absence of ongoing replication, and decrease in their localizations over time could be indicative of potential repair in non-replicating cells.

### Nucleotide Excision Repair generates ssDNA gaps for localization of replisome components in non-replicating cells

How do replisome components localize in non-replicating cells? SSB foci under these conditions indicates the presence of ssDNA stretches long enough to accommodate SSB tetramers (>30 nt) (Bell et al., 2015; Lohman & Ferrari, 1994). In replicating cells, ssDNA tracts are thought to be generated as a result of helicase activity that continues to unwind double-stranded DNA ahead of the replisome that has encountered a lesion (Belle et al., 2007). It is unclear how such tracts are formed in non-replicating cells. We wondered whether this could be mediated via pathways involved in DNA damage repair and tolerance. Given that several repair pathways are regulated under the SOS response (Baharoglu & Mazel, 2014), we first assessed the induction of the response in non-replicating cells under DNA damage. For this, we measured the induction of YFP from an SOS inducible promoter (P_*sidA*_) integrated on the *Caulobacter* chromosome at the *xyl* locus (Badrinarayanan et al., 2015) (Figure 3A). We found that non-replicating cells turned on the DNA damage response after MMC exposure, providing further evidence for the formation of ssDNA gaps in such conditions (Figure 3A). We thus asked whether the SOS response is essential for the formation of such gaps or if the activation of this response is a consequence of gap generation. Deletion of the SOS activator, *recA,* did not perturb localization of DnaN under damage. However, RecA was essential for dissociation during damage recovery as DnaN foci persisted in non-replicating cells lacking RecA (Figure 3B). These observations suggest that a RecA-independent pathway is required for regulating the association of replisome components with DNA in cells that are not undergoing active DNA synthesis.

**Figure 3:**
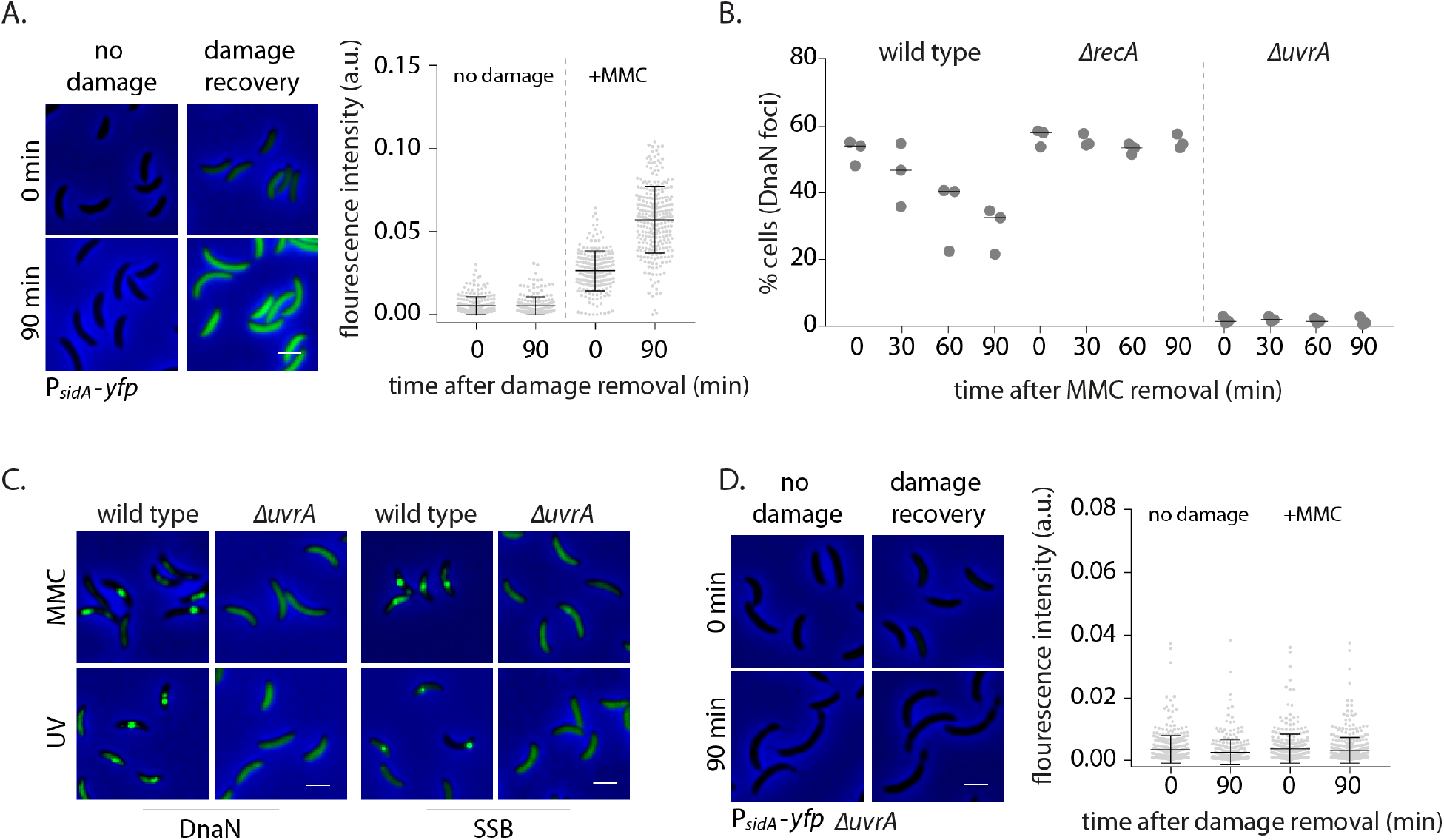
Nucleotide Excision Repair generates ssDNA gaps for localization of replisome components in non-replicating cells. (A) SOS induction is measured by assessing the expression of YFP from an SOS-inducible promoter (*P_sidA_-yfp*). On the left are representative images of cells expressing the reporter at 0 or 90 min after MMC removal and control cells (no damage). On the right total fluorescence intensity normalized to cell area is plotted for both time points for cells with or without damage treatment. Each dot represents a single cell. Mean and SD are shown in black (n ≥ 219). (B) Percentage wild type, *ΔrecA*, or *ΔuvrA* swarmer cells with DnaN foci 0, 30, 60 and 90 min after DNA damage recovery (n ≥ 308 cells, three independent repeats). (C) Representative images of wild type or *ΔuvrA* swarmer cells expressing SSB-YFP or DnaN-YFP, treated with MMC or UV. (D) As (a) for cells lacking *uvrA* (n ≥ 325).

In most organisms, helix distorting lesions are recognized and excised by Nucleotide Excision Repair (NER) (Kisker et al., 2013). Short gaps generated during this process could also be converted into longer stretches of ssDNA tracts under certain conditions as seen in eukaryotic systems (Sertic et al., 2011, 2018), thus requiring extensive DNA synthesis outside the active replication fork (Figure S3A). To test if this could be the mechanism by which replisome components associate with DNA in cells that are not replicating, we assessed the involvement of NER in orchestrating the same in *Caulobacter* swarmer cells. We observed that unlike wild type, non-replicating cells with deletion of *uvrA* (part of the NER pathway) did not form DnaN foci under MMC or UV damage (Figure 3B-C, Figure S3C). In contrast, percentage cells with DnaN foci in a *ΔmutL* background, deficient in mismatch repair (Marinus, 2012) was similar to wild type, indicating that mismatch repair did not contribute to loading of the β-clamp in non-replicating cells (Figure S3D).

Thus, our data suggest that lesion processing by NER alone results in the formation of ssDNA gaps on which replisome components can localize in non-replicating cells. Consistent with this, we observed lack of SSB localization in *ΔuvrA* cells both under MMC and UV damage (Figure 3C, Figure S3B and S3C). Furthermore, cells without NER were deficient in SOS induction (Figure 3D), suggesting that NER-mediated gap generation serves two functions: a. Providing ssDNA substrate for recruitment of SSB and other replisome components to these regions, b. Induction of the SOS response. Together, this facilitates ssDNA gap filling in non-replicating *Caulobacter*.

### SOS-induced low fidelity polymerase, DnaE2, is essential for subsequent dissociation of replisome components

As stated above, we observed that *ΔrecA* cells were not deficient in DnaN recruitment to ssDNA gaps. However, given that these cells had persistent β-clamp foci, we wondered what would be the requirement for RecA or the SOS response in ssDNA gap filling. We ruled out a role for homologous recombination in this process as our experimental setup of non-replicating swarmer cells (with a single chromosome) does not permit gap-filling by recombination, due to absence of a homologous template for repair (Figure 1A, bottom panel). In addition, we also conducted our damage recovery experiments in cells lacking the recombination protein RecN (Vickridge et al., 2017), an essential component of recombination-based repair in *Caulobacter* (Badrinarayanan et al., 2015). In this case too, we observed association, followed by dissociation of β-clamp foci as seen in case of wild type cells (Figure S4A).

Reports in *E. coli* as well as eukaryotic systems (including yeast and human cells) have suggested that ssDNA gaps generated by NER can sometimes be filled by specialized polymerases like Polκ (Janel-Bintz et al., 2017; Kozmin & Jinks-Robertson, 2013; Sertic et al., 2018). Given that the SOS response is activated in non-replicating cells (Figure 3A), it is possible that gap filling in *Caulobacter* swarmer cells is mediated via such specialized polymerases expressed under this regulon (Galhardo, 2005). Although we were unable to generate a functional fluorescent fusion to *Caulobacter* low-fidelity polymerase DnaE2, we confirmed that DnaE2 is expressed in our experimental conditions (Figure S4B) and that deletion of *dnaE2* resulted in severe sensitivity of a steady-state population of cells to MMC-treatment (Figure S2A, Figure S4F). To test the involvement of DnaE2 in gap filling, we conducted our damage recovery experiments in cells deleted for the same. Similar to *ΔrecA* cells, we found that non-replicating cells lacking *dnaE2* had persistent DnaN foci during damage recovery (Figure 4A-B). For example, in case of wild type, 52% cells had foci after 30 min of 0.5 μg/ml MMC treatment and this number reduced to 30% 90 min post MMC removal. In contrast, in case of *ΔdnaE2* cells, 61% cells had foci after 30 min of damage treatment and this number remained constant even after removal of MMC from the growth media. DnaN foci in *ΔdnaE2* cells was significantly higher than wild type after 90 min of damage recovery in case of UV damage as well, at the two doses of damage tested (Figure S4D).

**Figure 4:**
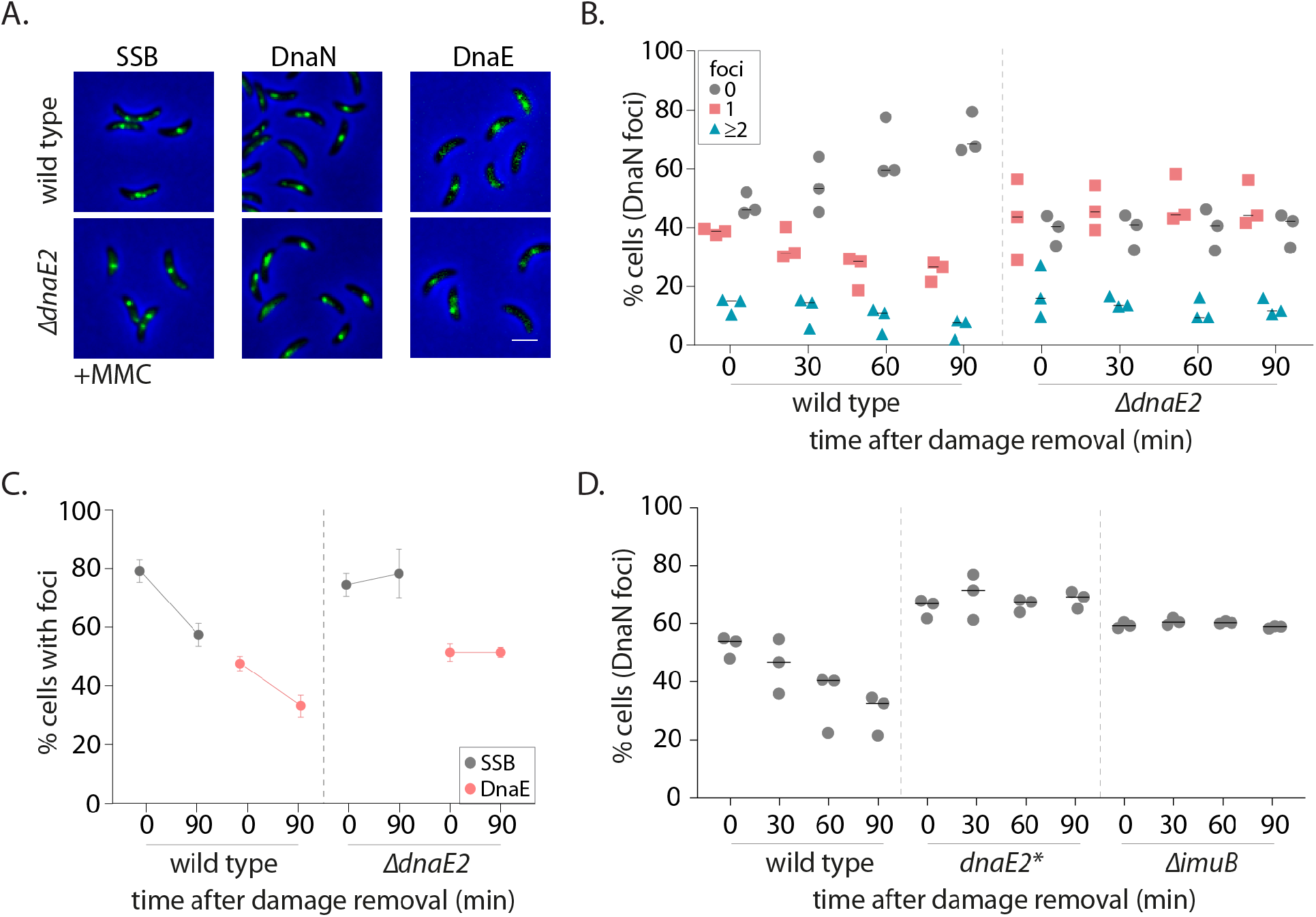
SOS-induced low fidelity polymerase, DnaE2, is essential for subsequent dissociation of replisome components. (A) Representative images of wild type or *ΔdnaE2* swarmer cells with SSB-YFP, DnaN-YFP or DnaE-YFP after MMC treatment. (B) Percentage wild type or *ΔdnaE2* swarmer cells with 0, 1, or ≥2 DnaN foci at 0, 30, 60 and 90 min of DNA damage recovery (n ≥ 467 cells, three independent repeats) (C) Percentage wild type or *ΔdnaE2* swarmer cells with SSB or DnaE foci at 0 and 90 min of DNA damage recovery (n ≥ 325 cells, mean and SD from three independent repeats). (D) Percentage wild type, *dnaE2* catalytic mutant (*dnaE2\**) or *ΔimuB* swarmer cells with DnaN foci at 0, 30, 60, and 90 min of MMC damage recovery (n ≥ 342 cells, three independent repeats. wild type data from Figure 2B).

Replisome persistence in the absence of *dnaE2* appeared to be a dose-dependent phenomenon (Figure S4C). At low dose of MMC treatment (0.125 μg/ml), fewer cells had DnaN foci post DNA damage exposure (14.5% cells). The number further reduced to 9.5% during recovery in a DnaE2-independent manner. However, the percentage of cells with persistent β-clamp foci increased with increasing concentrations of damage in the absence of *dnaE2,* with minimal recovery observed at 0.5 - 0.75 μg/ml of MMC treatment (Figure S4C). The following observations in our study lend additional support to the proposed idea that a specialized polymerase is required for gap filling ssDNA tracts generated by NER at higher doses of DNA damage: a. Persistence of components of PolIIIHE (DnaE and DnaN) in the absence of DnaE2. Apart from β-clamp foci, we found that the replicative polymerase, DnaE, was also unable to dissociate during damage recovery in cells lacking *dnaE2* (Figure 4C), suggesting that the replicative polymerase alone cannot complete synthesis across NER-generated ssDNA tracts. Such lack of dissociation after localization was found to be the case for SSB as well, again suggesting that ssDNA gaps persisted in the absence of DnaE2 (Figure 4C). b. Requirement for DnaE2-mediated synthesis. To test whether synthesis by DnaE2 contributes to gap filling in non-replicating cells, we mutated two residues known to be essential for DnaE-mediated synthesis in wild type (Lamers et al., 2006; Pritchard & McHenry, 1999). These residues have been mutated previously in *M. smegmatis* DnaE2, where it was shown to inhibit DnaE2-dependent mutagenesis (Warner et al., 2010) (Figure S4E). In case of *Caulobacter* as well, *dnaE2*\* showed similar growth defects as *ΔdnaE2* under MMC damage (Figure S4F). In our experimental regime, we found that cells expressing catalytically inactive DnaE2 also had persistent DnaN foci during damage recovery, as seen in the case of cells lacking the specialized polymerase (Figure 4D).

To assess contribution of DnaE2 to damage-induced mutagenesis, we conducted mutagenesis assays by measuring the frequency of rifampicin resistance generation in the population of cells subject to damage, with or without recovery in non-replicating conditions. We observed that this polymerase was responsible for all damage-induced mutagenesis in our experimental regimen (Figure S4G). However, the genetic complexity of this experiment and the confounding effects of replication during the out-growth period preclude us from conclusively interpreting if this mutagenesis mediated by DnaE2 occurs in non-replicating, replicating or both phases of the cell cycle.

Finally, we also tested the requirement for accessory protein ImuB in DnaE2 function. ImuB is an inactive Y-family polymerase and carries a β-clamp binding motif. It is thought to act as a bridge between DnaE2 and the clamp, likely facilitating DnaE2 binding to the clamp for function (Warner et al., 2010). In *Caulobacter,* it is co-operonic with DnaE2 and is also expressed in response to SOS activation (Galhardo, 2005). When we conducted our recovery experiments in cells lacking *imuB*, we observed that these cells also exhibited persistent DnaN foci, as seen for cells lacking *dnaE2* (Figure 4D). These results are consistent with the idea that DnaE2-mediated synthesis contributes to gap-filling and subsequent dissociation of replisome components in non-replicating cells.

### DnaE2 activity on NER-generated ssDNA gaps enhances survival of non-replicating cells under DNA damage

Taken together, our data provide *in vivo* support for cross-talk between two independent genome integrity maintenance systems (NER and specialized, low-fidelity polymerases) in non-replicating bacteria. What could be the relevance of this in the context of damage recovery and survival of bacteria that are not actively replicating? To investigate the impact of NER-mediated DnaE2 activity in *Caulobacter* swarmer cells, we assessed the growth dynamics of these cells once released into replication-permissive conditions with three parameters: a). Time to division and percentage cells with successful division events after release in replication permissive conditions (as a read-out for division restoration post DNA damage clearance) b). Cell length restoration (as a read-out for SOS deactivation following DNA damage clearance). c). Cell survival measured via viable cell count assays.

To measure division restoration, we released replication-blocked swarmer cells into media containing IPTG (to allow for replication initiation via induction of *dnaA*) either immediately after damage treatment or after 90 min of damage recovery. We followed single cells via time-lapse imaging to assess the time taken to first division after replication initiation (Figure 5A-B). Control cells without damage treatment and with/ without additional 90 min arrest in swarmer stage were able to robustly resume cell growth and division with >94% cells undergoing their first division within 240 min of release into replication-permissive conditions. Based on this, we followed cell division dynamics for cells treated with damage during this time window, wherein control cells (without damage) were successfully able to restore cell division. In MMC-treated conditions, we found that cells released into replication-permissive conditions immediately after damage treatment did not recover efficiently, with only 5% cells undergoing their first division within 240 min (Figure 5C). In contrast, wild type cells that were provided time for damage recovery before re-initiating replication, showed restoration of cell division in the same time period, with 30% cells undergoing at least one division and 9% cells undergoing ≥ 2 divisions within 240 min (Figure 5B-C). These recovery dynamics were dependent on DnaE2 as only 7% cells lacking *dnaE2* underwent divisions even when they were provided the same time duration as wild type for damage recovery before replication re-initiation (Figure 5B-C). Thus DnaE2-mediated gap filling provided a significant survival advantage to non-replicating cells as measured by their ability to robustly restore cell cycle progression and cell division.

**Figure 5:**
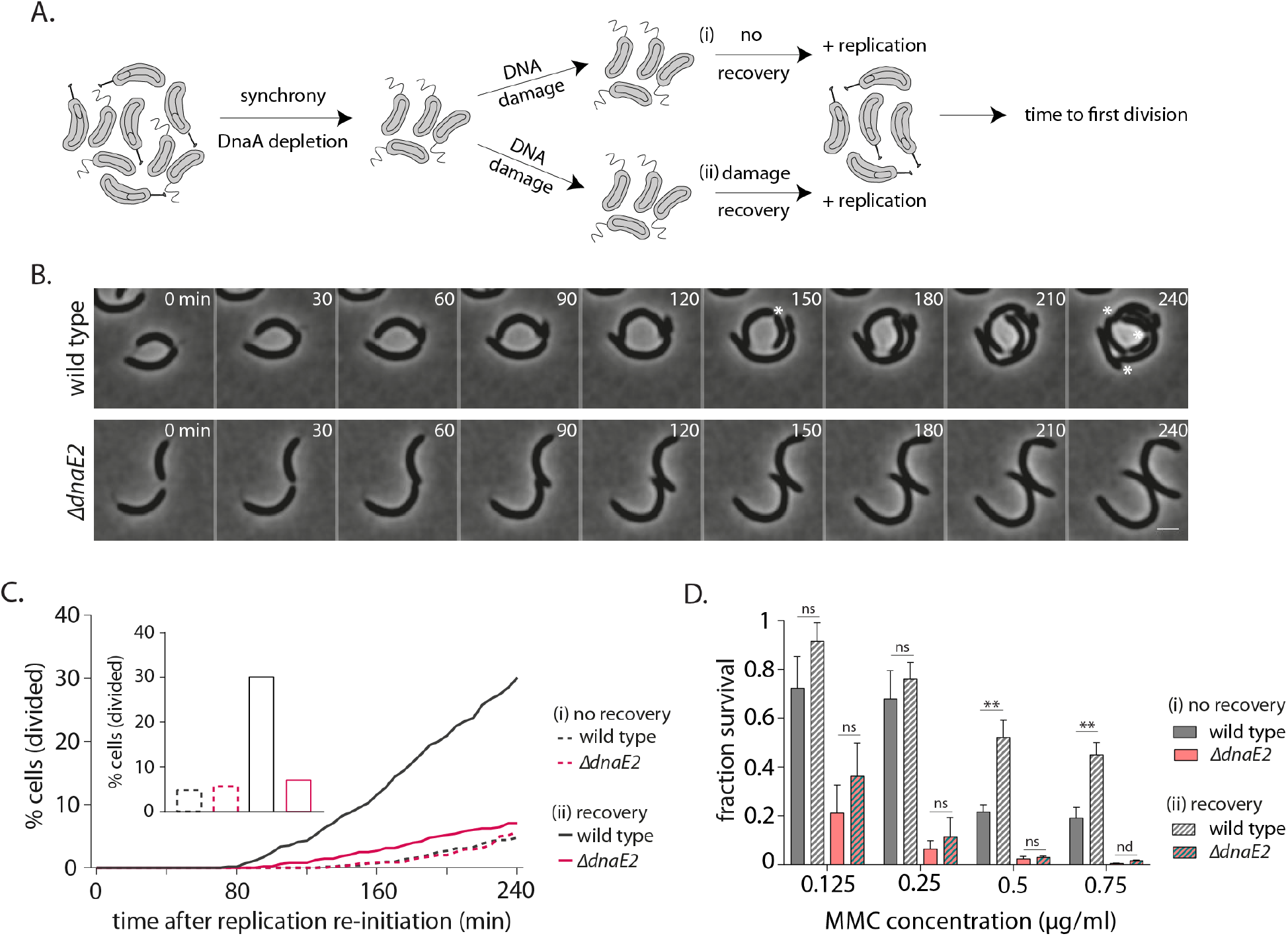
DnaE2 activity on NER-generated ssDNA gaps enhances survival of non-replicating cells under DNA damage. (A) Schematic of experimental setup used to assess the impact of lesion repair/ tolerance in non-replicating cells. After MMC treatment for 30 min, cells are either released into replication-permissive media ((i) no recovery) or allowed to grow for 90 min without damage and then released into replication-permissive media ((ii) damage recovery). Cells are followed via time-lapse microscopy and time to division is estimated. Control cells are taken through the same growth regimes, however, no damage is added to the culture. (B) Representative time-lapse montage of wild type or *ΔdnaE2* cells in replication-permissive media after DNA damage recovery. Cell divisions are marked with white asterisk. In the panel shown here three divisions were scored in wild type, while none were observed in *ΔdnaE2* cells (C) Percentage cell division over time after replication re-initiation for wild type and *ΔdnaE2* cells either without (i. no recovery) or with (ii. recovery) damage recovery time in replication-blocked conditions (n ≥ 368 cells). Percentage cells divided at 240 min in each of these conditions is summarized in the graph inset. (D) Survival of wild type and *ΔdnaE2* cells either without (i. no recovery) or with (ii. recovery) damage recovery time in replication-blocked conditions measured via estimation of viable cell count (three independent repeats). Fraction survival was calculated by normalizing viable cell count under DNA damage to that without DNA damage. Error bars represent mean with SD from three independent experiments.

To further assess the consequence of gap filling, we measured the cell length distributions for cells released into replication-permissive conditions with or without 90 min of DNA damage recovery (Figure S5A). Continued cell length elongation would be reflective of a continued division block, a hallmark of the SOS response. On the other hand, cell length restoration would be expected only for those cells where damage has been repaired. We found that cells that did not face damage (with or without *dnaE2*) had a median cell length of 4.6 μm after 90 min incubation in swarmer conditions. At 240 min after re-initiation of replication, the cell length distribution was restored close to a wild type-like pattern (control) with the median cell length dropping to 2.9 μm (Figure S5A, ‘no damage’). Length restoration was also observed in wild type cells able to engage in DnaE2 mediated gap filling in the 90 min recovery window (Figure S5A, ‘+ damage, recovery, wild type’). This restoration in cell length was dependent on the time provided for damage recovery as well as presence of DnaE2, as in both cases, cells continued to elongate after release into IPTG-containing media (Figure S5A, ‘+damage, no recovery’ and ‘+damage, recovery, *ΔdnaE2’*).

To lend support to these cell biological observations, we modified our recovery setup to measure viable cell counts instead (Figure S5B). For this we assessed the ‘fraction survival’ as defined by the viable cell count obtained for cultures with damage treatment and normalized to the viable cell count for cultures without damage treatment. We observed that wild type cells that were released into replication-permissive conditions without the 90 min window of damage recovery were significantly compromised in growth, with fraction survival reducing to 0.19 at higher doses of damage in the absence of recovery. On the other hand, in case of cells grown with the possibility of undergoing 90 min of damage recovery, the fraction survival increased to 0.45 at the highest dose of damage used (Figure 5D). We then asked whether the survival advantage observed during recovery was dependent on DnaE2 action. Consistent with a dose-dependent effect on replisome persistence in the absence of DnaE2, we also observed that DnaE2 had a significant impact on the replication-independent survival advantage at higher doses of DNA damage. As expected from a steady state population, we found that cells deleted for *dnaE2* were severely compromised for survival at all doses of damage used (Figure 5D). However, at higher doses of damage, cells lacking *dnaE2* had similar loss in viable cell counts whether or not they were given a 90 min window of recovery; only around 0.01 fraction survival was observed with or without damage recovery in case of cells lacking *dnaE2*, in contrast to the 0.45 fraction survival observed in case of wild type cells provided a period of damage recovery (Figure 5D). Thus, there was a significant component of enhanced survival in cells that could undergo repair in non-replicating conditions and this survival advantage was dependent on DnaE2.

In summary, our cell biological and genetic read-outs suggest that DnaE2-mediated gap-filling enables cell cycle restoration and cell division licensing when non-replicating cells are allowed to re-initiate DNA replication. In the absence of such recovery (either *dnaE2* deletion or cells grown without the window of recovery), cell division is compromised and cells continue to elongate, a hallmark of persistent DNA damage and hence continuously active SOS response. The impact of delayed cell division and subsequent cell length elongation is directly observed when viable cell count of the population is measured, with a dose-dependent effect on survival in cells compromised for recovery due to deletion of *dnaE2.*

## Discussion

DNA lesion repair and tolerance has been well-studied in a replication-centric paradigm (Gabbai et al., 2014; Indiani et al., 2005; Marians, 2018). Characterization of error-prone polymerases in *E. coli* has informed us about mechanisms of tolerance that could occur at the replication fork or behind it, in gaps generated due to replisome skipping over the lesion, followed by repriming downstream of it (Chang et al., 2019; Gabbai et al., 2014; Indiani et al., 2005). However, DNA damage is a universal event that can occur across all stages of the cell cycle, including in non-replicating conditions. This can have effects on transcription and could also perturb replication progression upon re-initiation (Jeiranian et al., 2013; Lang & Merrikh, 2018; Rudolph et al., 2007). For example, bacteria such as *Caulobacter* have distinct cell cycle phases including a non-replicating swarmer state, with a single copy of its chromosome. Hence it is imperative that DNA damage gets cleared out efficiently even in these conditions. Here we provide *in vivo* evidence for NER-coupled DnaE2 function that is active in non-replicating bacteria. This study complements a growing body of work that supports the possibility of low-fidelity polymerase-mediated synthesis (including mutagenesis) in replication-independent conditions (such as in stationary phase cells) across domains of life (Bull et al., 2001; Corzett et al., 2013; Janel-Bintz et al., 2017; Sung et al., 2003; Yeiser et al., 2002) and underscores the need to reconsider function of such polymerases outside canonical, isolated roles of lesion bypass during replication.

### DNA damage repair in non-replicating cells: requirement for DnaE2

Here, we develop a system to specifically assess mechanisms of damage repair and tolerance employed in cells that are not undergoing active DNA synthesis. Using replication initiation-inhibited *Caulobacter* swarmer cells, we show that lesions are dealt with in two main steps: a. damage processing by NER to reveal ssDNA gaps and b. gap filling by SOS-induced specialized polymerase, DnaE2. Due to absence of a second copy of the chromosome in our assay (all cells are non-replicating and have a single chromosome), role of homologous recombination in this process is unlikely. Hence, our observations are consistent with a scenario where the low-fidelity polymerase alone is sufficient to synthesize across ssDNA gaps generated by NER action. Why is there a need for a specialized polymerase during gap-filling of NER-generated substrates? We explore two possible scenarios here:

1. Conventionally NER is thought to generate gaps of approximately 12 nucleotides during lesion repair, which can be gap-filled by DNA PolI (Kisker et al., 2013). However, localization of SSB in our experiments suggests that gaps generated are >30 nucleotides, enabling SSB tetramerization and binding (Bell et al., 2015; Lohman & Ferrari, 1994). How are longer ssDNA tracts generated? Previous reports in *E. coli* as well as yeast and human cells have implicated a role for exonuclease activity in generating longer ssDNA tracts on NER substrates. In these studies, it was proposed that such activity would occur on problematic intermediates generated during NER activity, including closely-spaced opposing lesions that are generated under high doses of DNA damage (Janel-Bintz et al., 2017; Kozmin & Jinks-Robertson, 2013; Sertic et al., 2018). Indeed, our observations on lack of dissociation of replicative polymerase (PolIII) in the absence of DnaE2 as well as dose-dependent impact on cell survival would be consistent with a speculative model where NER-mediated excision results in the production of lesion-containing ssDNA that requires synthesis by a specialized polymerase.
2. It is equally plausible that DnaE2 contributes to gap filling independent of the presence or absence of a DNA lesion. Gap filling activity has been suggested previously for *E. coli* PolV and eukaryotic Polκ (Isogawa et al., 2018; Janel-Bintz et al., 2017; Ogi & Lehmann, 2006). Furthermore, recent studies on post-replicative gap-filling have proposed a scenario where long patches requiring synthesis are accessed by both replicative and TLS polymerases (PolIV and PolV) in *E. coli* (Isogawa et al., 2018). Thus, error-prone polymerases can function beyond their canonical role in replication-associated lesion bypass (Fujii & Fuchs, 2020). In case of non-rpelicating *Caulobacter* cells, it is possible that this polymerase can access the β-clamp and hence participate in gap-filling, given the observed increase in DnaE2 levels via SOS induction.

While our mutagenesis assays (measuring generation of rifampicin resistant mutations during damage) suggest that DnaE2 contributes to all MMC-induced mutagenesis, we were unable to satisfactorily disentangle the individual contributions from non-replicating vs replicating conditions (Figure S4G). Hence, we cannot reliably distinguish between the ‘gap-filling alone’ or ‘gap filling associated with lesion bypass’ activities of this polymerase in our present study. It must be noted though, that a role for DnaE2 in gap filling alone has not been reported before. In addition, unlike *E. coli,* it is the only polymerase implicated in TLS-associated functions (mutagenesis) in the bacteria that encode it. Thus, while we cannot provide a conclusive answer to this question, irrespective of the specific nature of DnaE2 activity, our work underscores a novel and necessary function for this highly conserved specialized polymerase in conjunction with NER in replication-independent conditions (discussed further below).

### ssDNA gaps generated by NER serve two functions

Previous studies in *E. coli* have found that NER activity in GGR is dependent on the activation of the SOS response (Crowley & Hanawalt, 1998). In contrast, our results suggest that NER functions upstream of the SOS response in non-replicating *Caulobacter*. Although *uvr* genes are SOS-induced even in *Caulobacter* (da Rocha et al., 2008), it is possible that basal levels of Uvr proteins are sufficient to carry out damage scanning and subsequent processing. Indeed, in *E. coli,* basal UvrA levels are variable, but range from 9 to 43 copies in minimal media to more than 120 copies in rich media (Ghodke et al., 2020, Stracy et al., 2016). Thus, ssDNA gaps generated by NER serve two purposes: a. Activation of the SOS response for specialized polymerase expression; it is likely that in case of *Caulobacter,* RecA is essential only for turning on the SOS regulon as DnaE2-mediated synthesis has been previously shown to function independent of RecA (Alves et al., 2017; Galhardo, 2005), unlike *E. coli* UmuDC (Goodman, 2014; Nohmi et al., 1988).

b. Providing substrate for SSB and PolIIIHE localization and specialized polymerase-mediated gap filling. SSB localization on ssDNA could further facilitate recruitment and loading of the PolIIIHE. While PolIII activity could directly contribute to gap filling (Isogawa et al., 2018; Sedgwick & Bridges, 1974; Soubry et al., 2019), it is also likely that it is the loading of the β-clamp that is essential for DnaE2 activity (Bunting et al., 2003; Chang et al., 2019; Fujii & Fuchs, 2004; Jérôme Wagner et al., 2009). Additionally, recent studies have highlighted a role for SSB as well in enriching the local pool of PolIV at a lesion, thus enabling polymerase switching (Chang et al., 2020). It would be interesting now to ask how additional components (such as ImuB and other accessory components to DnaE2) contribute to the loading of the ‘specialized replisome’ outside the realms of active replication and whether the properties of the ssDNA gaps generated may vary under different damaging conditions (UV vs MMC).

The lack of a significant percentage of cells with multiple replisome foci under damage would suggest that some repair or replisome components could be limiting, resulting in sequential repair or synthesis events. Alternatively, it is also possible that competition between SSB and RecA for binding ssDNA results in lesser SSB foci than the number of potential ssDNA tracts. The components involved in this process would be important in governing the number of patches that can be synthesized across at a given instance as well as the duration of a synthesis event. Indeed, distinct modes of action and nature of lesions induced by diverse damaging agents (Bargonetti et al., 2010; Chatterjee & Walker, 2017; Mitchell & Nairn, 1989) may contribute to some differences in the dynamics of replisome association/ dissociation observed here for MMC vs UV damage (Figure S4C & S4D). Finally, although discussed in the context of non-replicating cells, it is plausible that this mechanism can occur spatially and temporally disconnected from the active replication fork in replicating cells as well, in support of observations in *E. coli* that have reported localization of PolIIIHE as well as specialized polymerases away from the active replication fork (Henrikus et al., 2018; Soubry et al., 2019).

### Relevance of NER-mediated specialized polymerase activity in non-replicating cells

Our study provides comprehensive insights into a mechanism of lesion repair and gap filling in non-replicating bacteria, that relies on coordinated action between NER and low-fidelity polymerases. Our data suggests a method through which an error-prone polymerase, DnaE2, functions beyond replication forks, impinging on its implications in growth and survival of non-replicating cells. The experimental system in this study provides a novel tool to investigate these mechanisms as well as additional players further and assess impacts of lesion repair and tolerance in replication independent, but metabolically active conditions, where damage to DNA via molecules including ROS is possible (Gray et al., 2019; Manina & McKinney, 2013), such as *Caulobacter* cells in ‘swarmer’ state or other cells outside S phase of cell cycle.

The relevance of the process described here is highlighted by the survival advantage it confers in non-replicating cells. It is possible that NER-coupled DnaE2-mediated synthesis helps avoid the problems associated with persistent ssDNA gaps (due to NER activity itself) or DNA damage on the chromosome, (Jeiranian et al., 2013; Murli et al., 2000; Rudolph et al., 2007). In line with this, a recent study in human cells showed that coordinated action of NER along with Y-family polymerase, Polκ, and exonuclease, Exo1, was crucial for gap filling and hence prevention of UV-induced double-stranded breaks in non-S phase cells (Sertic et al., 2018). Such a role for specialized polymerases in gap-filling has also been observed in case of yeast cells (Kozmin & Jinks-Robertson, 2013; Sertic et al., 2011). More generally, this work highlights the possibility of coordinated activity of repair and tolerance pathways canonically studied as functioning independently. The universality of the NER-mediated error-prone polymerase function described here is underscored by its functionality in a diverse range of model systems, from bacteria to yeast and human cells (Janel-Bintz et al., 2017; Kozmin & Jinks-Robertson, 2013; Sertic et al., 2018), independent of the type or family of error-prone polymerase (DnaE2 in *Caulobacter* vs PolIV/ PolV in *E. coli*) employed during gap-filling.

## Materials and methods

### Bacterial strains and growth conditions

Bacterial strains, plasmids and primers used in the study are listed in Supplementary file 1 (Modell et al., 2014; Skerker et al., 2005; Thanbichler et al., 2007). Construction of plasmids and strains are detailed in the Supplementary file 1. Transductions were performed using ɸCR30 (Ely, 1991). *Caulobacter crescentus* cultures were grown at 30°C in PYE media (0.2% peptone, 0.1% yeast extract and 0.06% MgSO_4_) supplemented with appropriate concentrations of antibiotics, as required. While growing strains carrying *dnaA* under an IPTG-inducible promoter, liquid media was supplemented with 0.5 mM IPTG and solid media with 1 mM IPTG. Microscopy experiments were performed in minimal media containing 1X M2 salts (0.087% Na_2_HPO_4_, 0.53% KH_2_PO_4_, 0.05% NH_4_Cl) supplemented with 1% PYE, 0.2% glucose, 0.01 mM FeSO_4_ and 0.01 mM CaCl_2_.

Non-replicating swarmer cells were isolated using synchrony protocols described previously (Badrinarayanan et al., 2015). Briefly, cells were grown overnight in minimal media supplemented with IPTG. Cultures in log-phase were depleted for DnaA via washing off IPTG and allowing cells to grow in IPTG (-) conditions for one generation (~130 min). Following this, cultures were synchronized and OD_600_ of resulting swarmer cells was adjusted to 0.1, prior to treatment with DNA damage. In case of MMC damage, appropriate volume of 0.5 mg/ml MMC (AG Scientific, #M-2715) stock (prepared by resuspending in sterile water) was added into the culture and incubated at 30°C for 30 min. Damage was washed off by pelleting down cells at 8000 rpm for 4 min and resuspending in fresh media. For UV damage, cultures were transferred to a 90 mm petri plate and exposed to specific energy settings in a UV Stratalinker 1800 (STRATAGENE). During recovery (after UV and MMC damage) cells were incubated for 90 minutes at 30°C and 200 rpm. For strains expressing *SSB-YFP*, *SSB-GFP* or DnaN-YFP under P_*xyl*_, 0.3% xylose was added 1.5 h prior to imaging. Replication re-initiation after damage recovery was achieved by inducing cultures with 0.5 mM IPTG. DNA damage treatment used was either 0.5 μg/ml MMC (30 min) or 75 J/m^2^ UV for all experiments, unless otherwise specified.

For flow cytometry analysis, 300 μl of cultures were fixed in 700 μl of 70% chilled ethanol and stored at 4°C until further processing. These samples were treated with 2 μg/ml RNaseA in 50 mM sodium citrate for 4h at 50°C. DNA was stained with Sytox green nucleic acid stain (5 mM solution in DMSO from Thermo Fisher Scientific) and analyzed on a BD Accuri flow cytometer.

### Fluorescence microscopy and image analysis

For time course imaging, 1 ml aliquots of cultures were taken at specified time points, pelleted and resuspended in 100 μl of growth medium. Images were taken without damage treatment (no damage control), after 30 min of damage treatment (+ damage) and again at 0, 30, 60, and 90 min after removal of DNA damage (recovery). Controls were taken through the same treatment regime, but no damaging agent was added to growth media. 2 μl of cell suspension was spotted on 1% agarose pads (prepared in minimal medium) and imaged. For time lapse imaging 2 μl cell suspension was spotted on 1.5% GTG agarose (prepared in minimal medium), grown inside an OkoLab incubation chamber maintained at 30°C and imaged at specific intervals for the indicated period of time. For cell division tracking after replication re-initiation, cells were grown on 1.5% GTG agarose in growth medium containing with 1 mM IPTG.

Microscopy was performed on a wide-field epifluorescence microscope (Eclipse Ti-2E, Nikon) with a 63X oil immersion objective (plan apochromat objective with NA 1.41) and illumination from pE4000 light source (CoolLED). The microscope was equipped with a motorized XY stage and focus was maintained using Perfect Focusing System (Nikon). Image acquisitions were done with Hamamatsu Orca Flash 4.0 camera using NIS-elements software (version 5.1). Images were analysed using ImageJ as well as Microbetracker or Oufti in MatLab (Paintdakhi et al., 2016; Sliusarenko et al., 2011). Values for random positions within each cell and relative position of replisome foci were generated using custom-written MatLab scripts. Graphs were plotted in GraphPad Prism 7.

### Survival assay

For calculating viability of asynchronous steady state population under DNA damage, *Caulobacter* cultures were grown in PYE with 0.5 mM IPTG to O.D_600_ of 0.3. Serial dilutions were made in 10-fold increments and 6 μl of each dilution (10-1 to 10-8) were spotted on PYE agar containing 1 mM IPTG and appropriate amounts of MMC. Growth was quantified by multiplying dilution factor of the last visible spot with number of colonies on the last spot. Percentage survival for each strain was calculated by normalizing growth of that specific strain on different concentrations of MMC to that on media without DNA damage.

For assessing survival of non-replicating cells under DNA damage, swarmer cells (10 ml, OD_600_ - 0.1) were taken through one of the experimental regimes (with or without recovery in non-replicating phase) as mentioned in Figure S5B. At the end of the experiment, they were serially diluted and plated on PYE agar containing 1 mM IPTG and viability colony counts were taken after 48 hours. Fraction survival was calculated by normalizing viability of MMC treated cells to those taken through the exact same experimental regime, but without DNA damage treatment.

### Rifampicin resistance assay

Swarmer cells (10 ml, OD_600_ – 0.1) were taken through the same experimental conditions (with or without recovery) as mentioned above for survival experiments (Figure S5B). At the end of the experiment, the cultures were spun down, re-suspended in 10 ml PYE containing 0.5 mM IPTG and grown at 30°C overnight (approx. 20 h). These cultures were plated on PYE agar containing 0.5 mM IPTG and 100 μg/ml Rifampicin. Rif resistant colonies were counted 48 hours after plating, and mutation frequencies were calculated by normalizing to viable cell count of that specific culture.

### Western blotting

At specific time points of the experiment, 1.5 ml aliquots of 0.1 O.D_600_ cultures were pelleted down at 10000 rpm for 5 min, pellets were snap frozen in liquid nitrogen and stored at −80°C until further use. Pellets were resuspended in SDS sample buffer, and boiled at 95°C for 10 min. Equal amounts of lysates were loaded on 6% SDS-PAGE gel, resolved at 100 V and transferred to PVDF membrane (BIO-RAD, #1620177) in a wet electroblotting system. Non-specific binding to the membrane was blocked with 5% Blotting-Grade Blocker (BIO-RAD, #170-6404), followed by probing with 1:2000 dilution of monoclonal anti-flag antibody (Sigma, #F1804) and 1:5000 dilution of HRP-linked anti-mouse secondary antibody (Cell Signaling Technology, #7076S). The blots were visualized after incubation with SuperSignal™ West PICO PLUS Chemiluminescent Substrate (Thermo SCIENTIFIC, #34577) using an iBright FL1000 imager (ThermoFisher SCIENTIFIC).

## Acknowledgements

The authors acknowledge assistance from Prachi Shinde and Ramya Rajagopalan for preliminary experiments and NCBS Central Imaging and Flow Facility (CIFF) for flow cytometry usage. The authors thank Dr. Rodrigo Reyes-Lamothe, Dr. Stephan Uphoff, Dr. Tung Le and members of the AB lab for helpful discussions and feedback on the manuscript. This work was supported by fellowships from DBT (DBT-RA), DST-SERB (PDF/2018/001164) (AMJ) and CSIR (SD) as well as grants from DBT-IYBA, HFSP CDA (00051/ 2017-C) and intramural funding from NCBS-TIFR (AB).

## Author contributions

AMJ: conception of project, experimental design, generation of tools and reagents, execution of experiments, data analysis, writing of manuscript. SD: execution of experiments, and generation of tools and reagents. IS: generation of tools and reagents, and execution of experiments related to western blots. AB: conception of project, experimental design and writing of manuscript.

---

**Figure S1:**
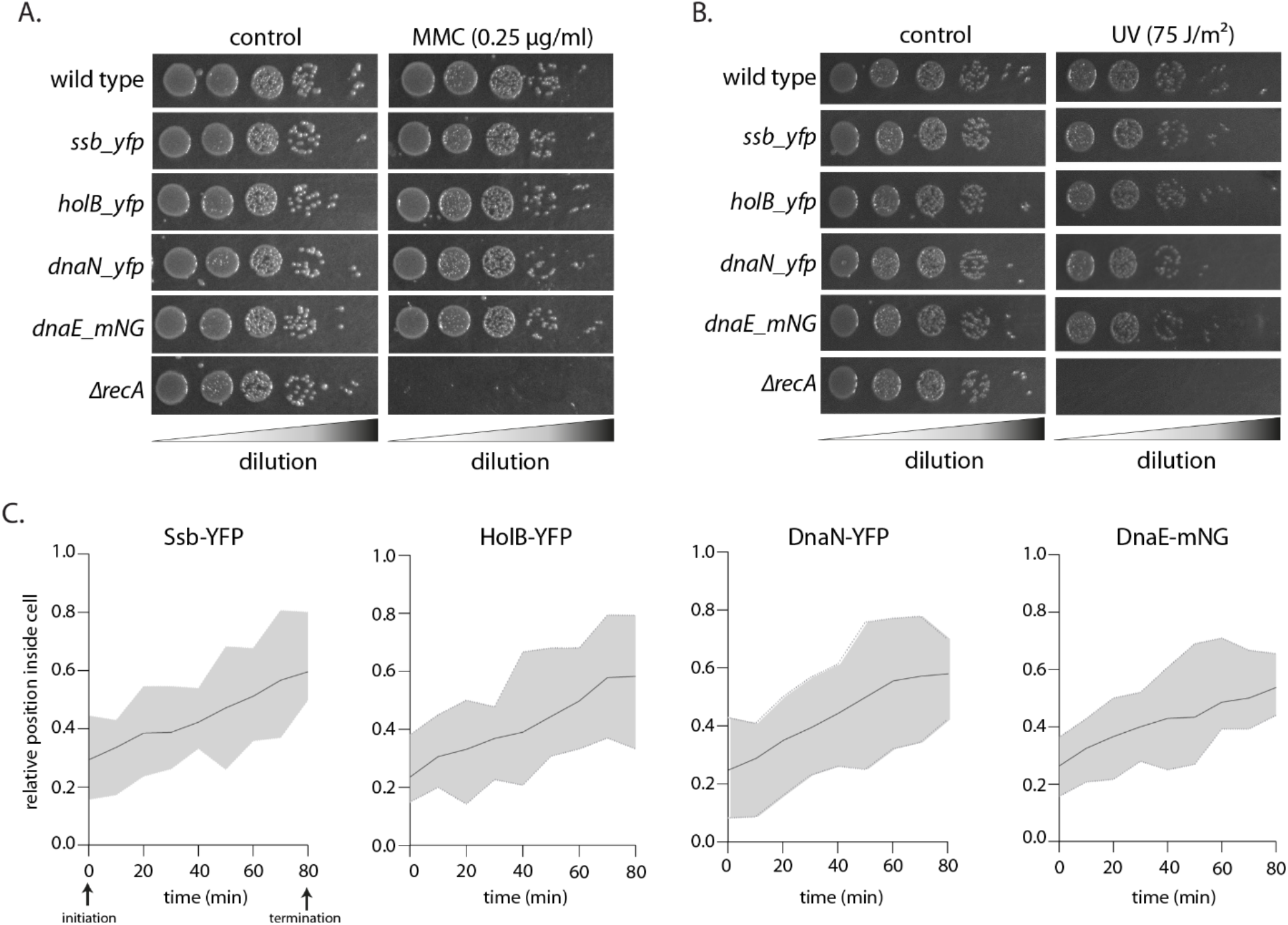
(A) Growth of fluorescently-tagged replisome strains with or without (control) MMC damage. For reference, growth of wild type (no tag) and *recA* deletion are also shown (representative image of one experiment from three independent repeats). (B) Growth of fluorescently-tagged replisome strains with or without (control) UV damage. For reference, growth of wild type (no tag) and *recA* deletion are also shown (representative image of one experiment from three independent repeats). (C) Relative position of fluorescently-tagged replisome components in *Caulobacter* cells during one round of replication (no damage induced). Localization of Ssb-YFP, HolB-YFP, DnaN-YFP or DnaE-mNG was tracked every 10 min using time-lapse imaging. A focus tended to localize at one cell pole at initiations and proceeded towards the opposite cell pole as replication progressed (n=25, solid line represents mean and shaded region depicts the upper and lower limit at specific time points).

**Figure S2:**
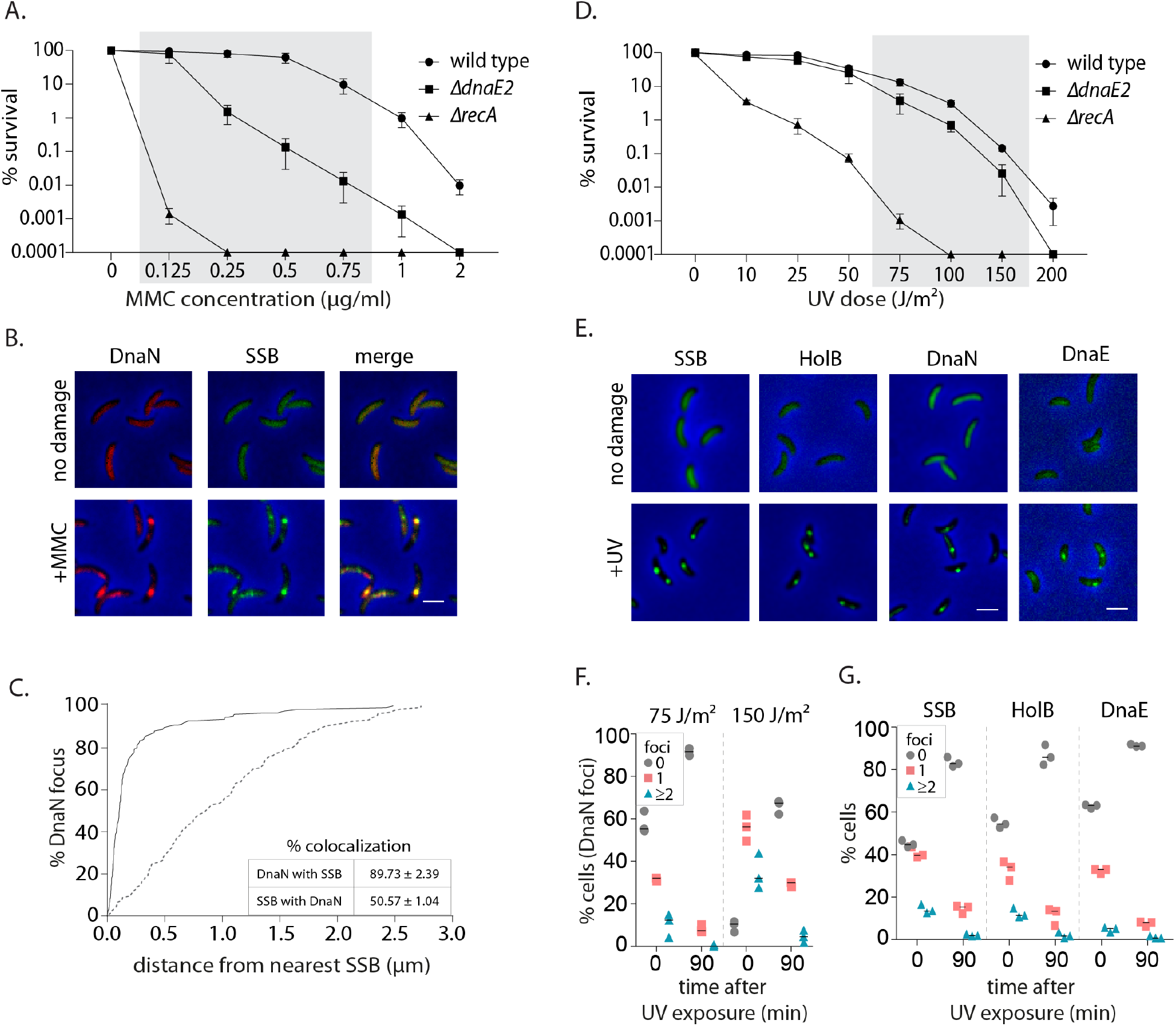
(A) Survival of wild type, *ΔdnaE2* and *ΔrecA* under different doses of MMC (mean and SD from three independent experiments). Shaded region depicts the concentrations used for experiments in this study. (B) Representative images of swarmer cells expressing DnaN-mCherry and SSB-GFP with or without MMC treatment (scale bar is 2 μm here and in all other images). (C) Distance of a DnaN focus from the nearest SSB focus is measured and cumulative frequency distribution is plotted (solid line). Dotted line is the distribution of distance between the DnaN focus and any random position inside the cell. In the inset, % colocalization for DnaN with SSB and vice versa is provided (mean and SD from three independent repeats). (D) Survival of wild type, *ΔdnaE2* and *ΔrecA* under different doses of UV (mean and SD from three independent experiments). Shaded region depicts the concentrations used for experiments in this study. (E) Representative images of swarmer cells expressing SSB-YFP, HolB-YFP, DnaN-YFP or DnaE-mNeonGreen with or without (no damage control) UV treatment. (F) Percentage wild type swarmer cells with 0, 1, or ≥2 foci of DnaN at 0 and 90 min after DNA damage recovery from 75 J/m^2^ or 150 J/m^2^ of UV (n ≥ 322 cells, three independent repeats). (G) Percentage wild type swarmer cells with 0, 1, or ≥2 foci of SSB, HolB or DnaE at 0 and 90 min after DNA damage recovery from 75 J/m^2^ of UV (n ≥ 334 cells, three independent repeats).

**Figure S3:**
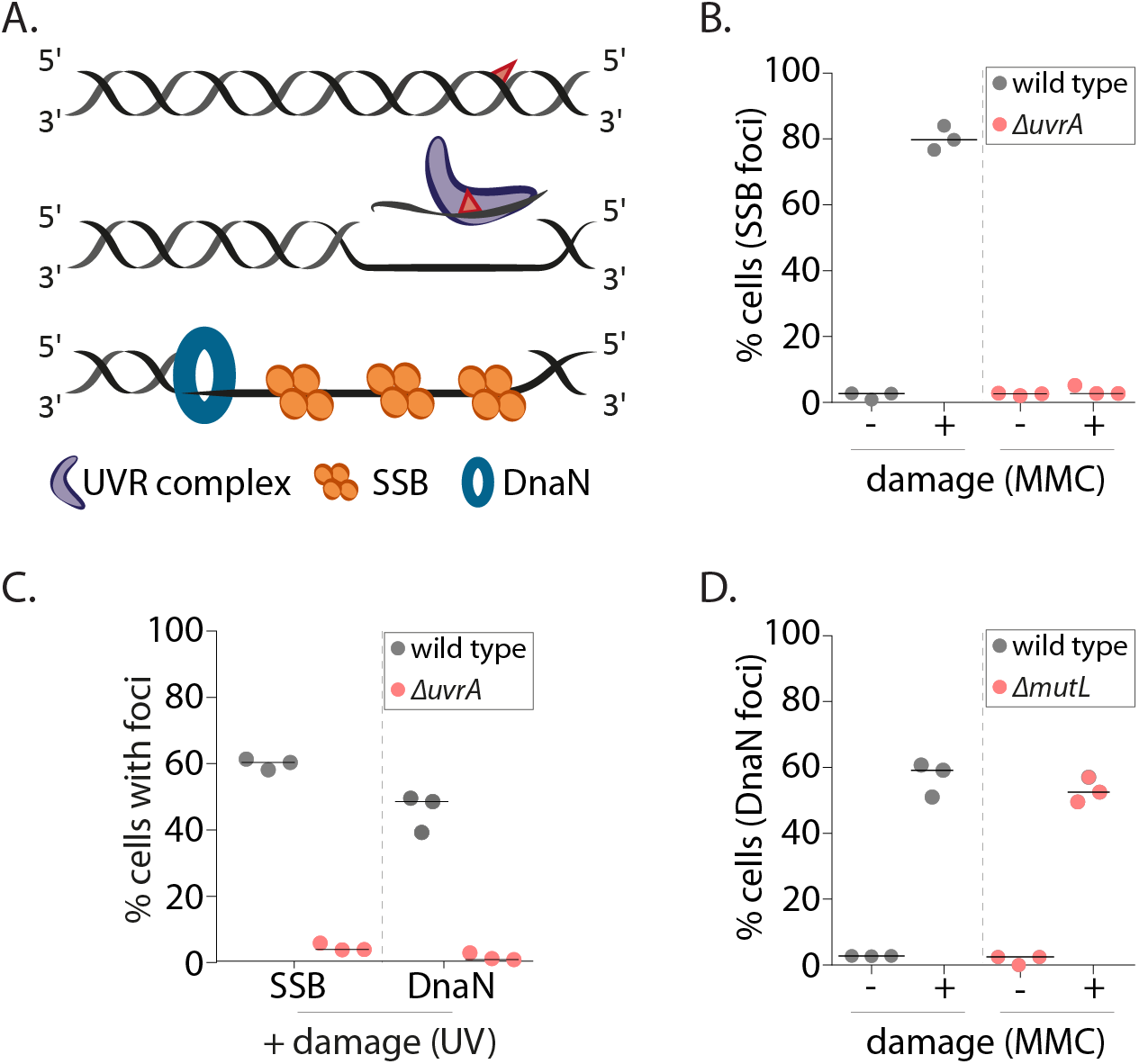
(A) Schematic of mechanism of single-stranded DNA gap generation by NER. (B) Percentage wild type or *ΔuvrA* swarmer cells with SSB foci with (+MMC) or without (-, control) damage treatment (n ≥ 325 cells, three independent repeats). (C) Percentage wild type or *ΔuvrA* swarmer cells with DnaN or SSB foci after DNA damage (UV) (n ≥ 340 cells, three independent repeats). (D) Percentage wild type or *ΔmutL* swarmer cells with DnaN foci with (+MMC) or without (-, control) damage treatment (n ≥ 324 cells, three independent repeats).

**Figure S4:**
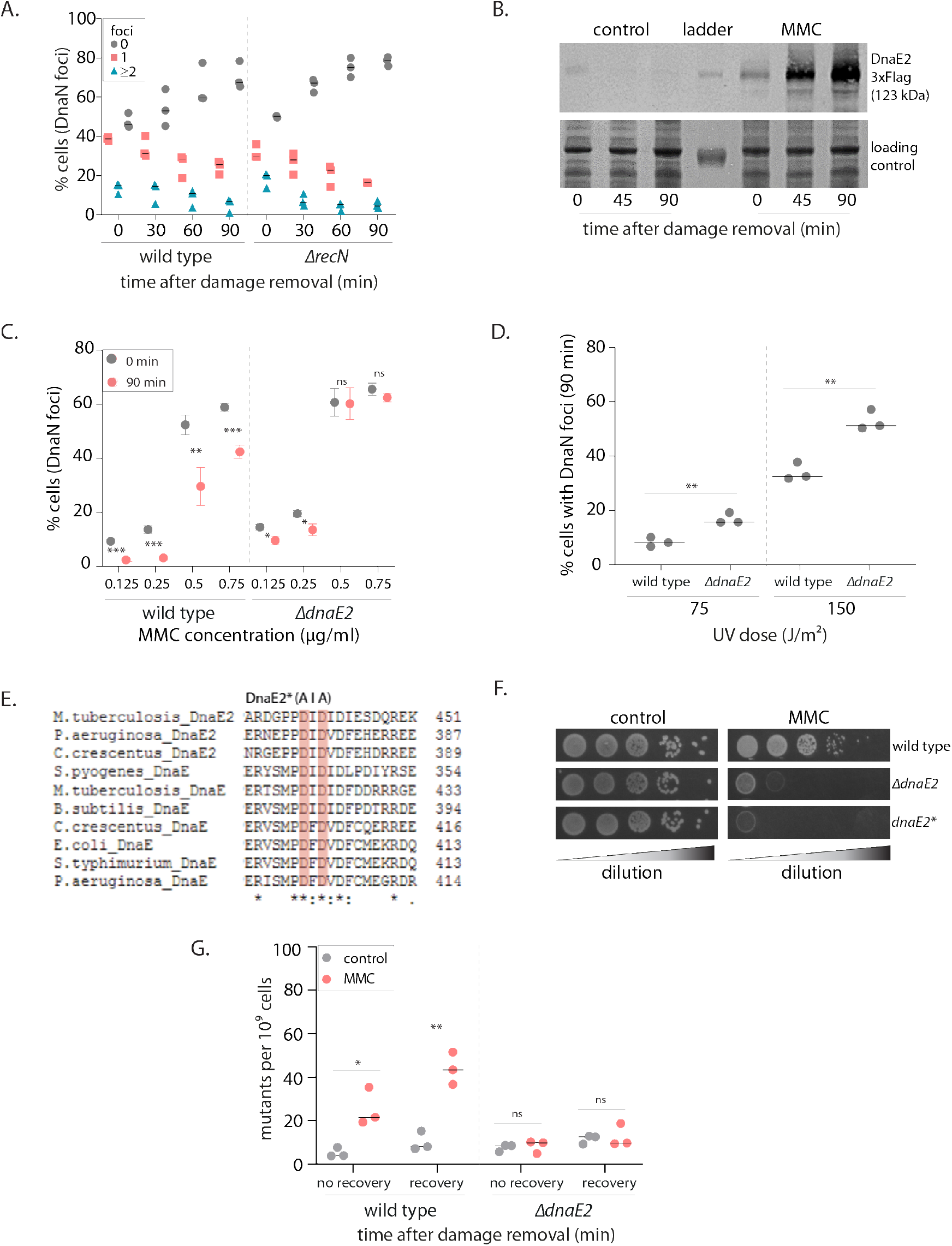
(A) Percentage wild type or *ΔrecN* swarmer cells with 0, 1, or ≥2 DnaN foci at 0, 30, 60 and 90 min of DNA damage recovery (n ≥ 309 cells, three independent repeats). (B) Representative image of a western blot on of DnaE2-3X-Flag during MMC damage recovery. As a control, cells without damage treatment are also probed for DnaE2 expression (image of one experiment from three independent repeats). (C) Percentage wild type or *ΔdnaE2* swarmer cells with DnaN foci at 0 and 90 min of DNA damage recovery (n ≥ 321 cells, mean and SD from three independent repeats, under indicated doses of DNA damage). Asterisks denote significant differences and ‘ns’ denotes not significant differences in unpaired t-tests here and in all other graphs. Specific p-values are summarized in Table 4. (D) Percentage wild type or *ΔdnaE2* swarmer cells with DnaN foci after 90 min of damage recovery post treatment with two doses of UV (n ≥ 332 cells, three independent repeats). (E) Multiple sequence alignment of a section of the catalytic domain of C-family polymerases from different bacteria. Conserved amino acid residues highlighted in pink have been mutated in DnaE2* (catalytic dead mutant) (Warner et al., 2010). (F) Growth of wild type, *ΔdnaE2* and *dnaE2\** strains with (MMC) or without (control) DNA damage (image of one experiment from three independent repeats). (G) Rifampicin resistant mutants that arise from wild type and *ΔdnaE2* cells treated with (MMC) or without (control) DNA damage. Cells were either immediately released into replication permissive media after damage removal (no recovery) or allowed to recover from damage for 90 min in non-replicating phase before release into replication permissive conditions (recovery). Dashed line shows median from three independent experiments.

**Figure S5:**
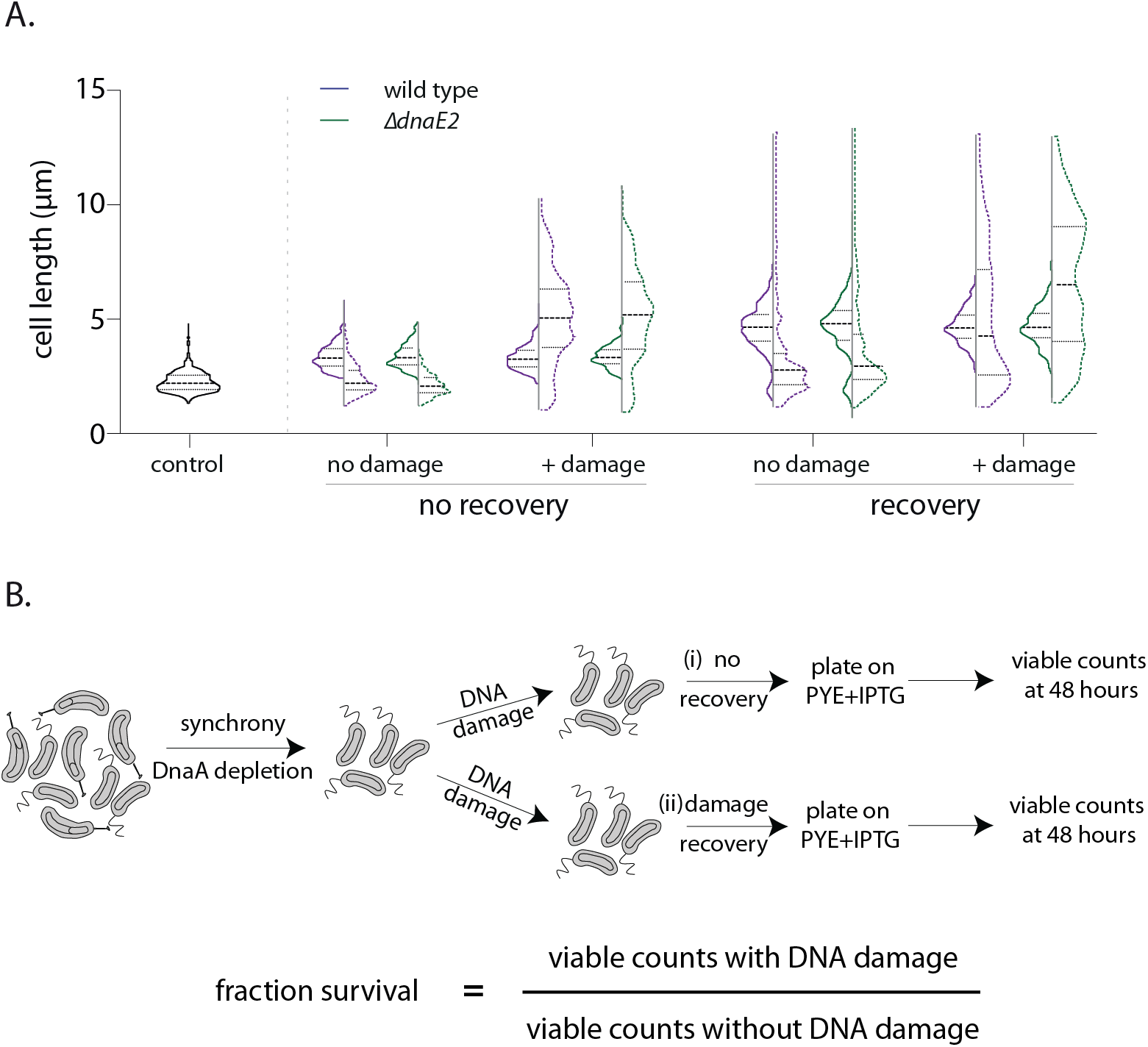
(A) Cell length distribution (with median) for wild type (purple) or *ΔdnaE2* (green) cells. Control cells were not treated with DNA damage, while + damage cells were exposed to MMC treatment for 30 min. Solid lines represent length distribution prior to release into replication-permissive conditions while dashed lines represent length distribution after 240 min in replication-permissive conditions. Median and inter-quartile range of the distribution is indicated. ‘No recovery’ and ‘recovery’ as outlined in Figure 5A (n ≥ 300 cells). (B) Schematic of experimental design to estimate survival advantage from recovery in non-replicating phase (Figure 5D). Fraction survival is calculated by normalizing viable cell counts obtained with damage to those obtained without damage. A similar experimental design was used for estimation of mutation frequencies (Figure S4G and materials and methods).

